# A new type of AmpC β-lactamases defined by PIB-1, a metal-dependent carbapenem-hydrolyzing β-lactamase, from *Pseudomonas aeruginosa*: structural and functional analysis

**DOI:** 10.1101/2024.03.05.583521

**Authors:** Francisco Javier Medrano, Sara Hernando-Amado, José Luis Martínez, Antonio Romero

## Abstract

Antibiotic resistance is one of most important health concerns nowadays. β-lactamases are the most important resistance determinants. Based on their structural and functional characteristics β-lactamases are grouped in four categories. AmpC β-lactamases are cephalosporinases presenting a set of highly conserved residues. Here we crystallized PIB-1, a *Pseudomonas aeruginosa* chromosomally-encoded β-lactamase. Its crystal structure shows it is an AmpC β-lactamase, although the number of conserved residues is low. Functional analysis showed that PIB-1 is able to degrade carbapenems but not the typical substrate of AmpC β-lactamases, cephalosporins. Besides, the catalytic activity of PIB-1 increases in the presence of metal ions. Metals do not bind to the active center and increase the degradation of the antibiotic. They induce the formation of trimers. This suggests that the oligomer is more active than the monomer. While PIB-1 is structurally an AmpC β-lactamase, the low sequence conservation, substrate profile and its metal-dependence, prompts us to position this enzyme as the founder of a new group inside the AmpC β-lactamases. Consequently, the diversity of AmpC β-lactamases might be wider than expected.

## Introduction

As recognized by several international agencies, antibiotic resistance constitutes one of most relevant problems for human health (Antimicrobial-Resistance-Collaborators, 2022). Knowing the mechanisms involved in the acquisition of resistance is critical for developing anti-resistance adjuvants (i.e. β-lactamase inhibitors) (Yahav *et al*, 2020) and evolution-based strategies to deal with the problem of antibiotic resistance (Sanz-García *et al*, 2023). Antibiotic inactivating enzymes constitute one of the most relevant mechanisms of resistance (Martinez & Baquero, 2014) and, among them, β-lactamases stand out as one of the most widespread group of antibiotic resistance determinants (Babic Hujer & Bonomo, 2006), being particularly relevant in the case of Gram-negative bacteria (Bush, 2018). Current classification of β-lactamases includes four groups, based on their functional and structural characteristics. Three of these groups (A, C and D β-lactamases) present serine in their active site, while the fourth (B-type β-lactamases) contains zinc ions in their active site, which is essential for their activity (Bush & Jacoby, 2010).

Several bacteria present chromosomally-encoded AmpC β-lactamases (Philippon *et al*, 2022), which are part of their intrinsic resistome (Fajardo *et al*, 2008). Besides, the genes encoding these enzymes can be present in mobile genetic elements (Bush & Bradford, 2020), which constitute an important risk for their spread (Martínez Coque & Baquero, 2015). As a consequence, AmpC β-lactamases is the broadest category of β-lactamases, with nearly 5000 members so far described (Naas *et al*, 2017). Despite their structural similarities, AmpC β-lactamases present a large variability (≤20% sequence identity). However, they present three well conserved catalytic motifs (64SXSK, 150YXN, 315KTG) and, in addition, 70 amino acid residues of these enzymes are also highly conserved (≥ 90%) (Philippon *et al*., 2022). The substrates of AmpC β-lactamases are mainly cephalosporins, although they can hydrolyze other β-lactams and, with much less efficacy, carbapenems (Philippon *et al*., 2022). However, some of them present affinity for carbapenems, which has allowed to solve the structure of AmpC β-lactamases binding the carbapenem imipenem in their catalytic domain (Beadle & Shoichet, 2002). Despite the binding of carbapenems, their hydrolysis rate is typically very low and, at least in occasions, AmpC enzymes have a “trapping effect” for carbapenems (Philippon *et al*., 2022). This binding, but poor hydrolysis, should not increase resistance to these carbapenems unless over-production of this enzyme occurs, potentially blocking or inactivating the total activity of the drug. Further, it has been stated that, carbapenems could also act as inhibitors of the cephalosporinase activity of AmpC β-lactamases (Papp-Wallace *et al*, 2014).

We previously described a *Pseudomonas aeruginosa* chromosomally-encoded β-lactamase that does not fit well in any of the above mentioned β-lactamases categories. The enzyme PIB-1 (from *Pseudomonas* imipenem β-lactamase) hydrolyzes imipenem but no other β-lactams, cephalosporins included. Based in this activity, PIB-1 would fit into the B group of the Ambler classification. However, based on a phylogenetic analysis using its primary sequence, it seems to be more closely related to AmpC β-lactamases, despite its substrate profile is uncommon in this group (Philippon *et al*., 2022). These results made us hypothesize that PIB-1 could define a new type of AmpC β-lactamases. Therefore, in this work we solved the structure of PIB-1 in the presence and in the absence of the carbapenem meropenem. Also, we have characterized the binding of metals to PIB-1, and its effects on the enzymatic activity of the protein. Our results prompt us to propose that the diversity of AmpC β-lactamases might be wider than expected.

## Results

### Structure of PIB-1

The β-lactamase PIB-1 was crystalized and its structure was determined in the presence and in the absence of the carbapenem meropenem. Three different crystal forms of PIB-1 were obtained. Two of them belonged to the space group P2_1_2_1_2_1_, apo state (8RLJ) and in complex with meropenem (8RLK), and the third one belongs to the space group R3 (8RLL), Se-Met derivatized protein in its apo state. The asymmetric unit of the orthorhombic crystal form contains six monomers (Fig 1A) and the R3 contains four monomers (Fig 1B). The alignment of monomer A from the three structures (apo state, Se-Met derivatized and in complex with meropenem) shows small differences among them (Fig 1C). The root mean square deviation (RMSD) was calculated, using LSQKAB (Kabsch, 1976) implemented in the CCP4 suite (Winn *et al*, 2011), for all atoms in the 385 residues being 0.428 Å between the complex with meropenem and the apo form in the space group P2_1_2_1_2_1_, 0.369 Å between the complex with meropenem and the Se-Met derivatized protein, and 0.406 Å between the apo form in the space group P2_1_2_1_2_1_ and the Se-Met derivatized protein. Between the amino acids that form the binding pocket, there are small differences in the position of the side chains for some of them, most likely due to the absence of substrate in the apo state (Fig 1C).

**Fig. 1.**
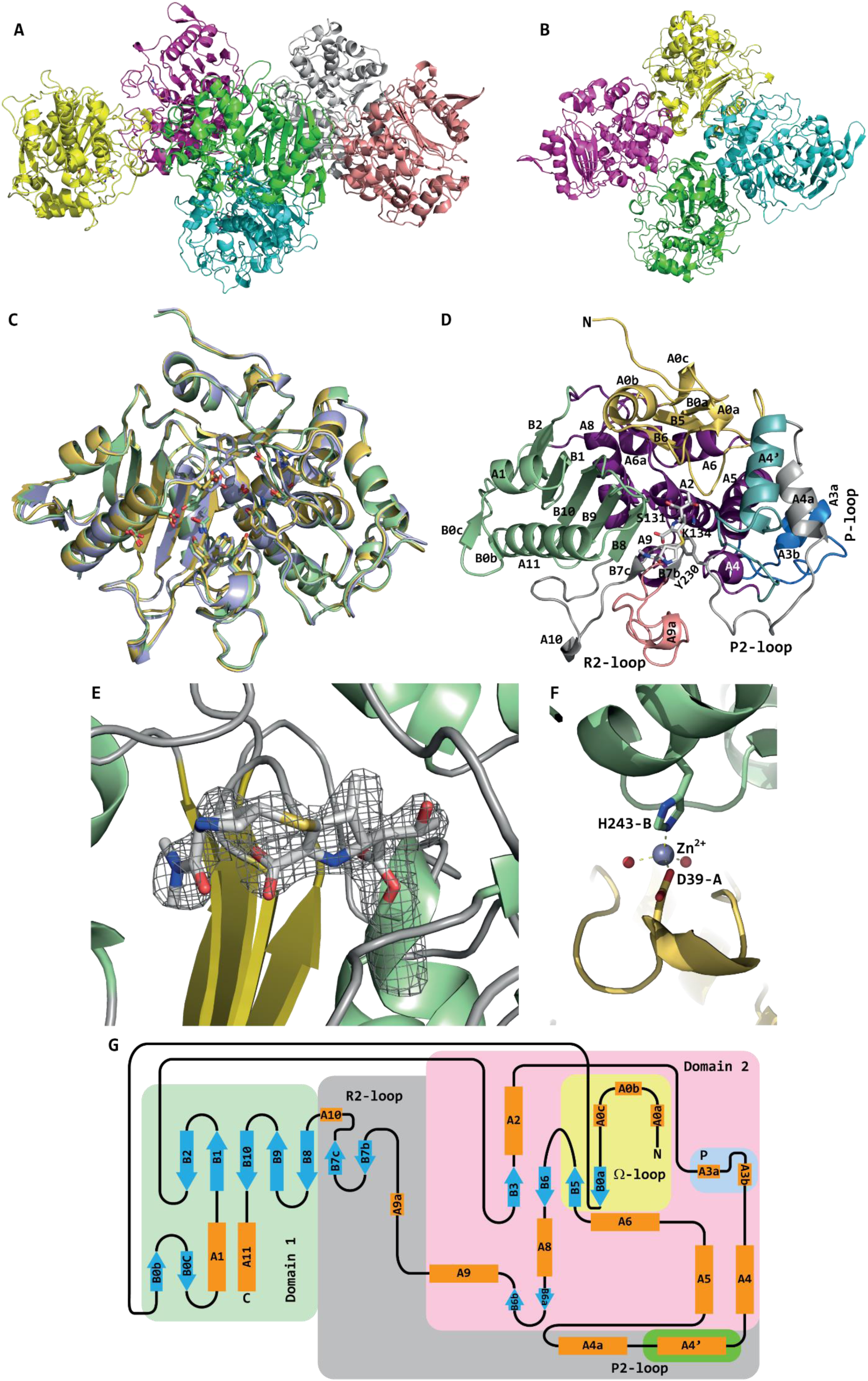
Structure and topology diagram of PIB-1. **A** Arrangement of the 6 monomers present in the crystal form with P2_1_2_1_2_1_ space group. **B** Arrangement of the 4 monomers present in the crystal form with R3 space group. **C** Structural alignment of the monomers of the apo form of PIB-1 in space group P2_1_2_1_2_1_ (light green), the Se-Met derivatized form in space group R3 (yellow orange) and the complex with meropenem in space group P2_1_2_1_2_1_ (light blue). **D** Structure of PIB-1 in complex with meropenem. The two major structural domains are colored in light green (domain 1) and deep purple (domain 2). The Ω-loop (yellow), the P-loop (blue), the R2-loop (salmon) and the P2-loop are also indicated. The three catalytic residues (Ser131, Lys134 and Tyr230) and the antibiotic are shown as sticks. **E** Electron density map of hydrolyzed meropenem bound to the active site of PIB-1 contoured at 1 σ. **F** Location of one Zn^2+^ atom bound to Asp39 from monomer A and His243 from monomer B and two waters. **G** Topology diagram for PIB-1. The two structural domains (domain 1 in light green and domain 2 in pink) are shown as colored background boxes. Helices are colored orange and strands are colored light blue. The Ω-loop and P-loop are indicated by yellow and light blue boxes, respectively. The P2 and R2 loops are additional elements, with respect to class A and D β-lactamases, located inside the gray box. An additional α-helix (A4’) in the P2 loop is indicated in the green box. The numbering scheme for the secondary structure elements follows that used by Bhattacharya *et al*. (Bhattacharya *et al*, 2014).

The structure of PIB-1 in complex with meropenem and its topology diagram are shown in Fig 1D-1G. In the crystal structure of PIB-1 in complex with meropenem, the antibiotic is only present in four of the six monomers. The electron density around the hydrolyzed antibiotic is shown in Fig 1E. In the structures of the Se-Met derivatized protein and the complex with meropenem only the first two residues of the sequence are not visible in the structure. The structure of the apo form has a greater number of residues not defined in the structure. The crystal structure of the protein in complex with meropenem presents three magnesium atoms that come from the crystallization solution, and seven zinc atoms that come from the protein solution. One of the zinc atoms is coordinated by two residues of the protein, side chains of Asp39 from monomer A and His243 from monomer B, and by two solvent molecules (Fig 1F). The other zinc atoms interact with the same residues from different monomers. The magnesium atoms are coordinated by six solvent molecules that do not directly contact with the protein. In one of these atoms, four of the solvent molecules interact with the main chain of residues Gly76 and Val77 from monomer A, and the main and side chain of Gln146 from monomer E (S1 Fig).

The mature form of PIB-1 has 387 amino acids, similar to the number of amino acids that present the AmpC β-lactamases (around 360 aa) and being longer than the class A and D β-lactamases, that are between 90 and 110 aa shorter than the typical AmpC β-lactamases (Bhattacharya *et al*., 2014) (around 130 aa shorter than PIB-1). The structure of PIB-1 has the two extra loops (P2 and R2) present in the AmpC β-lactamases (Fig 1D and 1G). Although with some notable differences (see below), the structure of PIB-1 resembles that of classical AmpC β-lactamases. In addition, meropenem is bound to the catalytic domain that contains the conserved motif SXXK which carries the catalytic serine. These structural characteristics support that PIB-1 is an AmpC β-lactamase, although it is capable of binding carbapenems in its catalytic domain.

### Binding pocket

The active site with bound hydrolyzed meropenem of PIB-1 is shown in Fig 2. The binding pocket for the substrate is made up by 19 residues. Residues Ser131, Lys134, Tyr230, Tyr334, Trp347 and Tyr338 are located at the bottom of the pocket; residues Tyr41, Phe130 and Ser283 form the back of the pocket; residues Glu187, Arg196 and Thr232 close the right side of the pocket; residues Gln367, Gly368, Ile369, Asp396 and Glu399 close the left side of the pocket; and residue Glu394 is placed on top of the substrate. The hydrolyzed antibiotic is covalently bound to the catalytic Ser131, and makes three hydrogen bonds between three residues of the protein and two oxygen atoms of the antibiotic (Fig 2, gray dashed lines).

**Fig 2.**
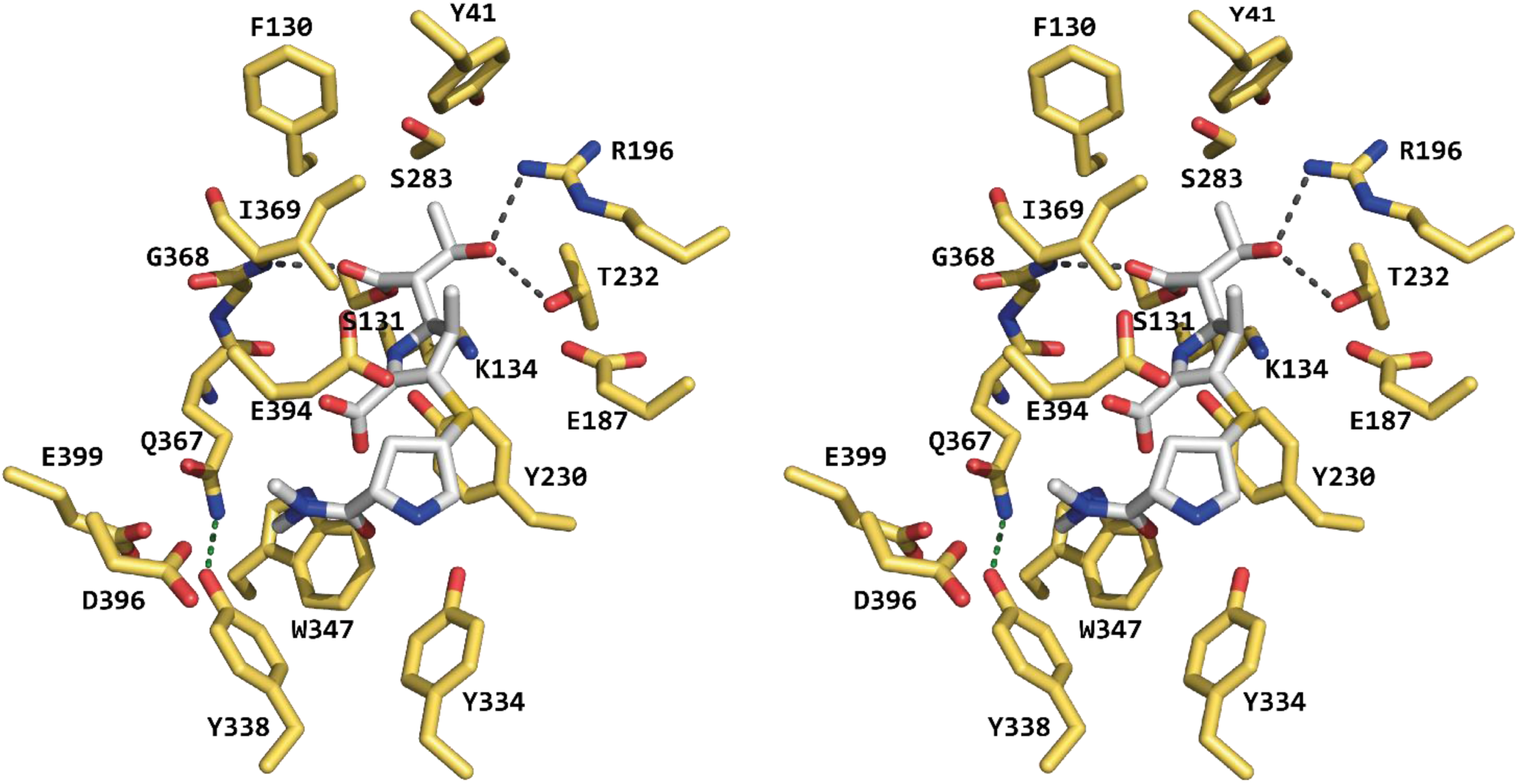
Binding pocket of PIB-1. Wall-eyed stereo pair of the residues forming the active site of PIB-1. The antibiotic meropenem is shown in gray and the protein residues in yellow. The dark-gray dashed lines represent hydrogen bonds between the protein and the antibiotic, and the green dashed line represents a hydrogen bond between protein residues.

### Comparison of PIB-1 with other AmpC β-lactamases

As above stated, the AmpC family of β-lactamases is very diverse with a highly variable primary structure, with levels of sequence identity as low as 20% (Brandt *et al*, 2017; Philippon *et al*., 2022). However, the structural alignment of different members of the family present in the PDB shows the extreme similarity among these proteins at the structural level. These members present a relatively high similarity, those from non-related organism range from 55-60% in similarity and those from related organism range from 80-98% similarity. These high similarity of the members of these family with solved structure has a high bias towards pathogenic organism. To get more insight regarding the similarity of PIB-1 with other AmpC β-lactamases, it was compared with PDC-1 from *P. aeruginosa* (Lahiri *et al*, 2014) (Fig 3). Due to the high structural similarity among the members structurally solved of this family, this comparison applies to all the members. Notably, the structural alignment of PIB-1 and PDC-1 shows clear differences. These differences mainly affect loops (P2, R2 and Ω-loop) that contribute to the substrate binding pocket (Fig 3A and 3B), a feature that could explain the substrate profile differences between PIB-1 and other AmpC β-lactamases (see below). The first, and most remarkable, difference is the extension of approximately 80 residues at the N-terminal that runs around the protein ending on the back of the substrate binding pocket (Fig 3A and 3B). The first 34 residues of this extension occupy the same region as residues 215 to 245 of PDC-1, but with a different conformation. It has three small helices and one β-strand that participates in the Ω-loop (A0a, A0b, A0c and B0a), then it runs around the back of the protein up to the location of the N-terminal of PDC-1, where it forms a β-hairpin (B0b and B0c) before joining helix A1. The second difference affects the P2-loop (PIB-1 residues 193 to 205) and represents an insertion in PIB-1 with respect to the typical members of this family (S3 Fig). It forms a long loop followed by an α-helix (A4’) and it contributes with two residues, Glu187 and Arg196, to the substrate binding pocket. Arg196 makes a hydrogen bond with the substrate meropenem (Fig 2). PDC-1 present a smaller loop in this region (residues 144 to 152) and occupies a smaller volume (Fig 1C, R2-loop). The third difference affects the R2-loop (PIB-1 residues 320 to 361) with an insertion of 12 amino acids (Supplementary Fig 3). Some of these amino acids (Tyr334, Tyr338 and Trp347) are involved in the binding of the substrate. This region, which contains less residues in PDC-1 (residues 284 to 337), presents a different conformation, presenting some loops and two short helices (Fig 3A and 3C).

**Fig 3.**
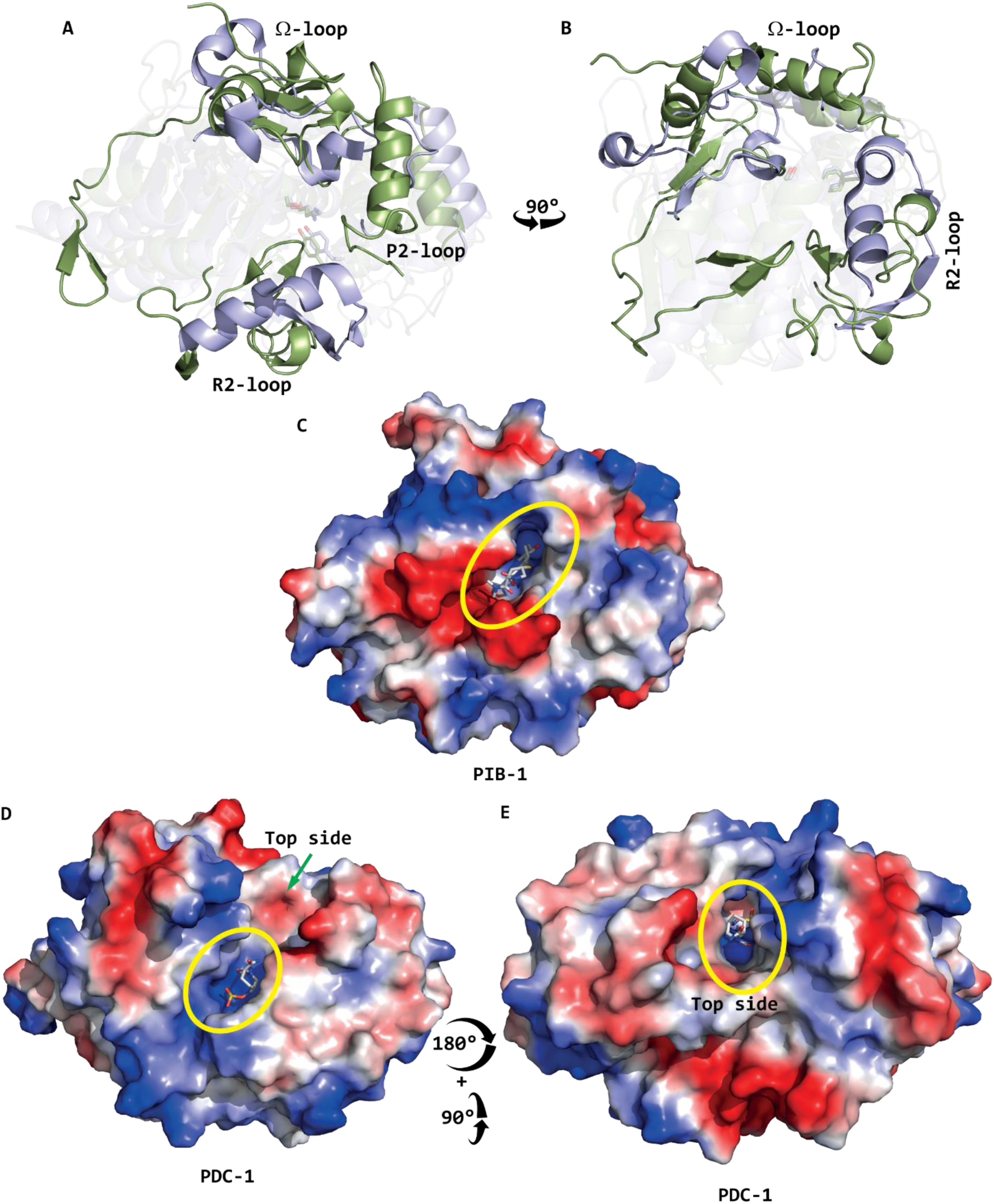
Structural comparison and electrostatic surface of PIB-1 and PDC-1. **A** Structural alignment of PIB-1 (light green) and PDC-1 from *P. aeruginosa* (light blue, PDB ID: 4OOY). The main differences affecting the loops P2, R2 and Ω are highlighted, and the rest of the protein is shown as transparency. **B** Rotation of 90° of the structural alignment. **C** PIB-1 from *P. aeruginosa* with meropenem bound to the active site. **C** PDC-1 from *P. aeruginosa* with the inhibitor avibactam bound to the active site. **D** PDC-1 from *P. aeruginosa* view from the top side. Blue represents negatively charged residues, red positively charged residues and white hydrophobic residues. The substrate binding pocket is indicated as a yellow ellipse. Substrate (PIB-1) or inhibitor (PDC-1) are shown as sticks inside the active site.

All these differences are reflected in the shape of the substrate binding pocket (Fig 3C-3E). This pocket is located in the middle of the enzyme (Fig 3C, yellow ellipse) as a deep cavity. It presents a positively charged region where the catalytic serine is located, and a large negatively charged region at the bottom of the pocket (Fig 3C). These shape limits the size of the possible substrates to those small enough to fit into this cavity. The substrate binding pocket in PDC-1 is also a deep cavity but it is open at the top side (Fig 3D and 3E; yellow ellipses). These shape limits the length of the substituents that the substrates can have to one side to be accommodated properly in the binding pocket, but the fact that it is open at the top allows these proteins to accommodate substrates with big substituents that will extend towards the top side of the binding pocket. Also, the character of the binding pocket in PDC-1 is markedly negative (Fig 3D and 3E). The different shape and electrostatic character of these binding pockets will, most likely, affect the type of substrate that these proteins are able to accommodate and process.

### PIB-1 is an atypical AmpC β-lactamase

It has been shown that, despite their diversity, the AmpC family of β-lactamases presents three conserved catalytic motifs and several conserved amino-acids(Philippon *et al*., 2022). In order to determine the presence of these elements in PIB-1, a sequence alignment, based on structural elements, was performed (Fig 4). The three conserved catalytic motifs typical of the Ambler class C β-lactamases can be observed in this alignment (Fig 4, yellow boxes). Among them, the only one clearly present in PIB-1 is the first one (SXXK), which carries the catalytic serine. The second motif does not present the typical YXN sequence. Instead, it has the sequence YST, which preserves just the Tyr, a member of the catalytic triad. Also, the third motif does not present the typical KSG sequence. It consists in an AQG sequence, only conserving the Gly at the third position. This Gly establish a hydrogen bond with meropenem (Fig 4).

**Fig 4.**
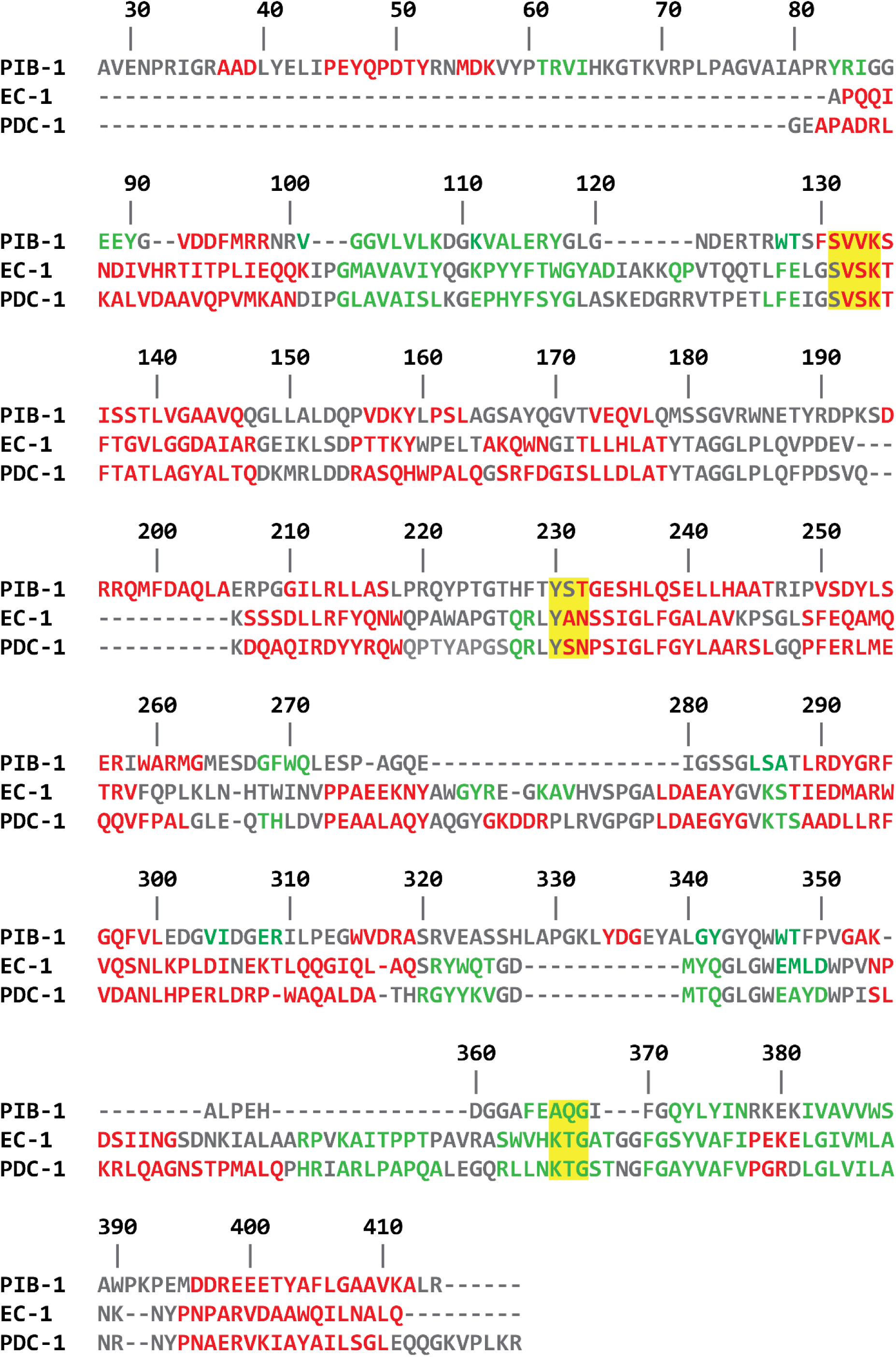
Sequence alignment of PIB-1 and of two AmpC enzymes, EC-1 from *E. coli* and PDC-1 from *P. aeruginosa*. The sequences were first aligned with CLUSTALW (https://www.genome.jp/tools-bin/clustalw) and further refined using the secondary structure elements. Residues in α-helices are colored red, in β-sheets green and in loops gray. The yellow boxes indicate the catalytic conserved motifs of AmpC β-lactamases. The numbering corresponds to the amino acid sequence of PIB-1.

Besides the three highly conserved catalytic motifs present in the AmpC β-lactamases, there are 70 amino acids that are also highly conserved. These includes 37 residues strictly conserved (100%) and 33 residues highly conserved (90-97%) in a data set of 32 β-lactamases (Mack *et al*, 2020; Philippon *et al*., 2022). Noteworthy, only 14 of the 37 strictly conserved residues are conserved in PIB-1 (S2 Table) and there is other residue with a conserved character of the residue (F→W). In addition, only 8 among the 33 highly conserved residues are present in PIB-1 (S2 Table). It is also worth mentioning that PIB-1 presents an extension of approximately 80 residues at the N-terminal, as well as four significant (more than three residues) deletions and two insertions. These findings indicate that PIB-1 presents some of the structural motifs that are conserved in AmpC β-lactamases, being particularly relevant the SXXK motif that contains the catalytic serine, but lacks several conserved amino acids from the other two catalytic conserved motifs. Together with the structure of PIB-1 above discussed, this indicates that, although PIB-1 can be classified into the Ambler class C of β-lactamases, a feature also supported by previous phylogenetic analyses (Fajardo *et al*, 2014), it forms a new branch within the family, where no β-lactamase had been included to date.

To delve deeper into this possibility, a phylogenetic analysis of PIB-1 and the different class C β-lactamases from the β-Lactamase DataBase (http://bldb.eu/Enzymes.php) (Naas *et al*., 2017) was performed (Fig 5). This database contains around 5000 AmpC β-lactamases, but it is highly biased in favor of bacterial pathogens, as *E. coli* of *P. aeruginosa*. Furthermore, each of the enzyme types in which the AmpC group is divided mainly contain alleles of the same type of β-lactamase presenting few changes among them. Consequently, to avoid potential biases in the phylogenetic analyses due to the over-representation of some β-lactamases, a representative member of each enzyme type was chosen for the analysis. As shown in Fig 5, PIB-1 forms an outgroup within the family of AmpC β-lactamases, further reinforcing that it defines a new uncharacterized branch in this family of proteins.

**Fig 5.**
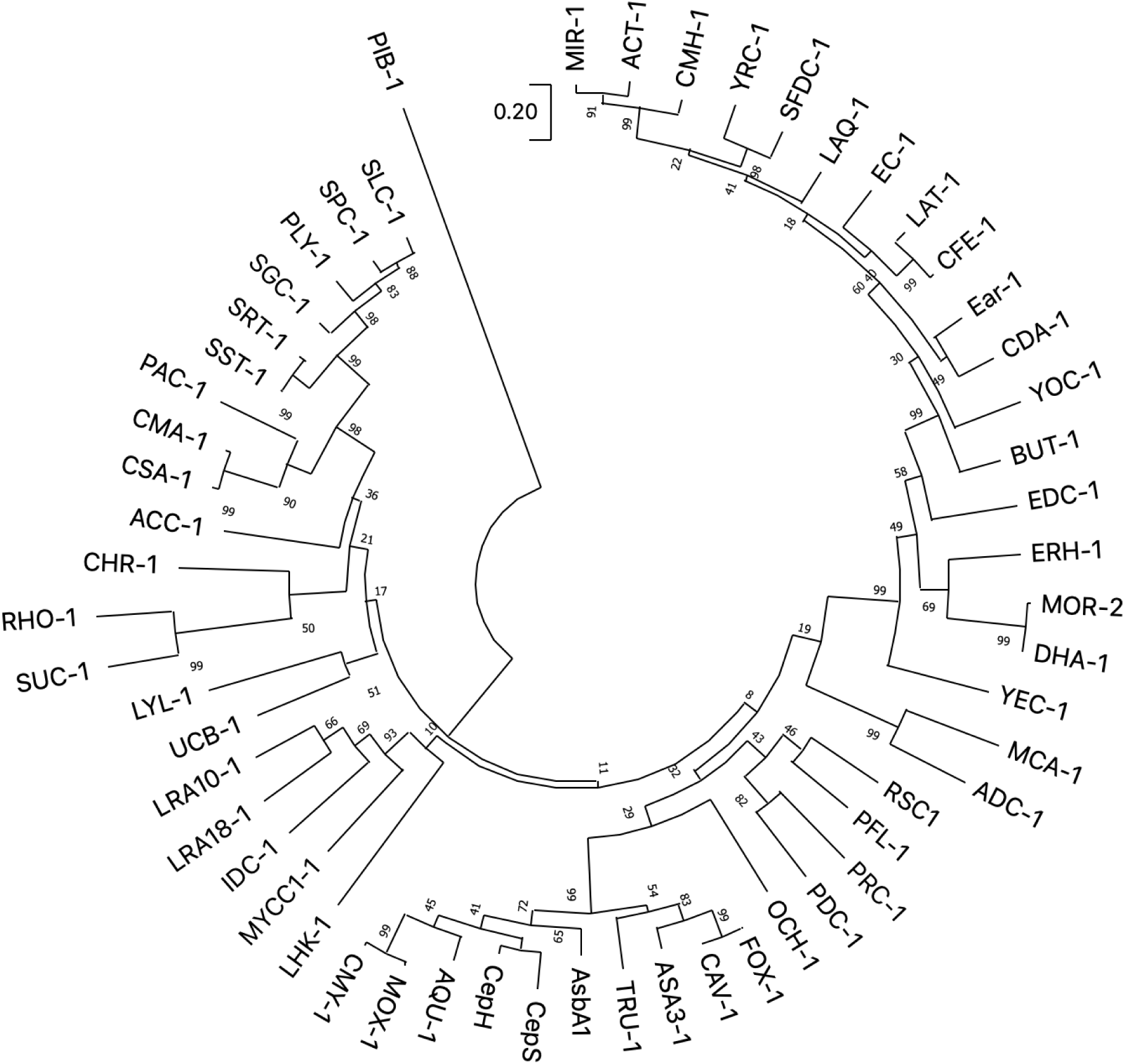
Phylogenetic analysis of PIB-1. The analysis was performed using the prototypes of the 55 class C β-lactamases present in the β-Lactamase DataBase (Naas *et al*., 2017). The analysis was performed with MEGA, using the Maximum Likelihood method and JTT matrix-based model (Tamura Stecher & Kumar, 2021). As shown, PIB-1 forms an outgroup within the AmpC family.

### Functional analysis of PIB-1

We previously showed that PIB-1 hydrolyzes imipenem but that it does not have an effect on the classical AmpC substrates, cephalosporins (Fajardo *et al*., 2014). Here we expanded the analysis to more β-lactam types and also to β-lactamase inhibitors, including avibactam, a well-known inhibitor of AmpC β-lactamases. To get functional information on the PIB-1 substrate profile, we measure the minimal inhibitory concentrations (MICs) in an *E. coli* strain (AF27) overexpressing PIB-1 or in an *E. coli* AF26 with the same plasmid without PIB-1 as a control, and without the intrinsic AmpC β-lactamase gene (Fajardo *et al*., 2014). In this experiment any variation in the level of susceptibility is solely attributable to the activity of PIB-1. MIC variations above 2-fold (analyzed using E-test strips, that discriminate slight MIC variations) indicated biologically relevant changes of resistance level, as previously described (Hernando-Amado *et al*, 2022a; Hernando-Amado *et al*, 2022b). As shown in Table 1, and in agreement with previous findings (Fajardo *et al*., 2014), the expression of PIB-1 reduces the susceptibility of *E. coli* to carbapenems, but does not have any effect in susceptibility to other β-lactams. Furthermore, the reduced susceptibility to imipenem of the strain expressing PIB-1 is not affected by avibactam, an inhibitor of AmpC β-lactamases, neither by clavulanic acid.

**Table 1.** Effect of PIB-1 expression on the susceptibility of *E. coli* to different β-lactams in the presence of 500 μM Zn^2+^.

| β-lactam family | Antibiotic | MIC (mg/L) | |
| --- | --- | --- | --- |
|  |  | AF26<sup>a</sup> | AF27<sup>b</sup> |
| Cephalosporins | Ceftazidime | 0.094 | 0.094 |
|  | Ceftazidime-avibactam* | 0.094 | 0.094 |
|  | Cefotaxime | 0.016 | 0.016 |
|  | Cefepime | 0.023 | 0.032 |
|  | Cefuroxime | 1.5 | 1.5 |
|  | Cefoxitin | 1.5 | 1.5 |
|  | Cefaclor | 0.5 | 0.75 |
|  | Ceftriaxone | 0.032 | 0.032 |
|  | Cefpirome | 0.047 | 0.047 |
| Penicillins | Amoxicillin | 1.5 | 2 |
|  | Amoxicillin-clavulanic acid** | 1.5 | 2 |
|  | Piperacillin | 0.75 | 0.75 |
|  | Piperacillin-tazobactam* | 0.75 | 0.75 |
|  | Ampicillin | 1.5 | 1.5 |
|  | Ampicillin-sulbactam* | 0.75 | 1.5 |
| Carbapenems | Imipenem | 0.125 | 0.75 |
|  | Imipenem-avibactam* | 0.125 | 0.75 |
|  | Imipenem-clavulanic acid** | 0.125 | 0.5 |
|  | Meropenem | 0.008 | 0.047 |
|  | Ertapenem | 0.064 | 0.19 |
| Monobactams | Aztreonam | 0.064 | 0.047 |

\* Avibactam, tazobactam and sulbactam were added to the plate at a concentration of 4 mg/L.
\*\* Clavulanic acid was added to the plate at a concentration of 2 mg/L.
<sup>a</sup>AF26, *E. coli* MI1443 strain carrying pUCP24 vector.
<sup>b</sup>AF27, *E. coli* MI1443 strain expressing PIB-1 cloned in the pUCP24 vector.

As mentioned, cephalosporins are the main substrates of AmpC β-lactamases but these enzymes can also degrade, with low efficacy, carbapenems (Jousset *et al*, 2019; Kaczmarek *et al*, 2006). In occasions, AmpC β-lactamases may be trapping carbapenems, something that in combination with a reduced permeability of the antibiotic associated with mutations in the porin OprD, could provide resistance to carbapenems (Philippon *et al*., 2022). Noteworthy, it has been recently described that mutations that increase the activity of *P. aeruginosa* against cephalosporins, decrease the capability of the enzyme for degrading carbapenems (Cabot *et al*, 2023), indicating an inverse correlation between carbapenems and cephalosporins degradation. PIB-1 would have suffered similar changes during evolution, increasing its activity against carbapenems and losing its capacity to hydrolyze cephalosporins.

### Oligomerization state

The analysis of the structure with PISA (Krissinel & Henrick, 2007) (https://www.ebi.ac.uk/pdbe/pisa/) predicts that the protein might be a trimer in solution in the presence of Zn^2+^, and it does not predict any oligomer in the Se-Met derivatized crystal. In order to clarify the oligomerization state of the protein we carried out SEC-MALS, analytical ultracentrifugation (sedimentation velocity) and gel filtration chromatography experiments (Fig 6). The SEC-MALS experiments shows the presence of monomers, dimers and trimers in solution and, in the presence of 500 μM Zn^2+^, there is only one species that corresponds to the trimer (Fig 6A). Sedimentation velocity experiments shows three species in the absence of metals (Fig 6B, gray line). In the presence of Zn^2+^, at concentrations around or below the K_d_ for the binding of the metal, the three species still co-exist, although, with the equilibrium shifted towards the trimer (Fig 6B, cyan and yellow lines). At concentrations higher than twice the K_d_ (120 and 500 μM), there are no monomers or dimers detectable, only trimers (Fig 6B, purple and green lines). High concentrations of Ni^2+^ (200 μM, Fig 6B, dark yellow line) or Co^2+^ (500 μM, Fig 6B, blue line) also shifts the equilibrium towards the trimer, but there still were small proportions of monomers and dimers. The molecular weights of the different species are summarized in S3 Table. Also, gel filtration chromatography was performed, indicating the presence of monomers, dimers and trimers in the absence of metals and at concentrations of Zn^2+^ up to 120 μM (Fig 6C; gray, cyan and yellow lines). At concentrations above 120 μM (Fig 6C, purple and green lines) there are only trimers in solution. Ni^2+^ and Co^2+^ reduced the proportion of monomers and dimers, but, differently from what happens with Zn^2+^, at the highest concentration analyzed (500 μM) they do not disappear completely (S2 Fig). The most likely arrangement for the trimer contains three Zn^2+^ atoms interacting with two monomers each of them to form a tight oligomer (Fig 6D).

**Fig 6.**
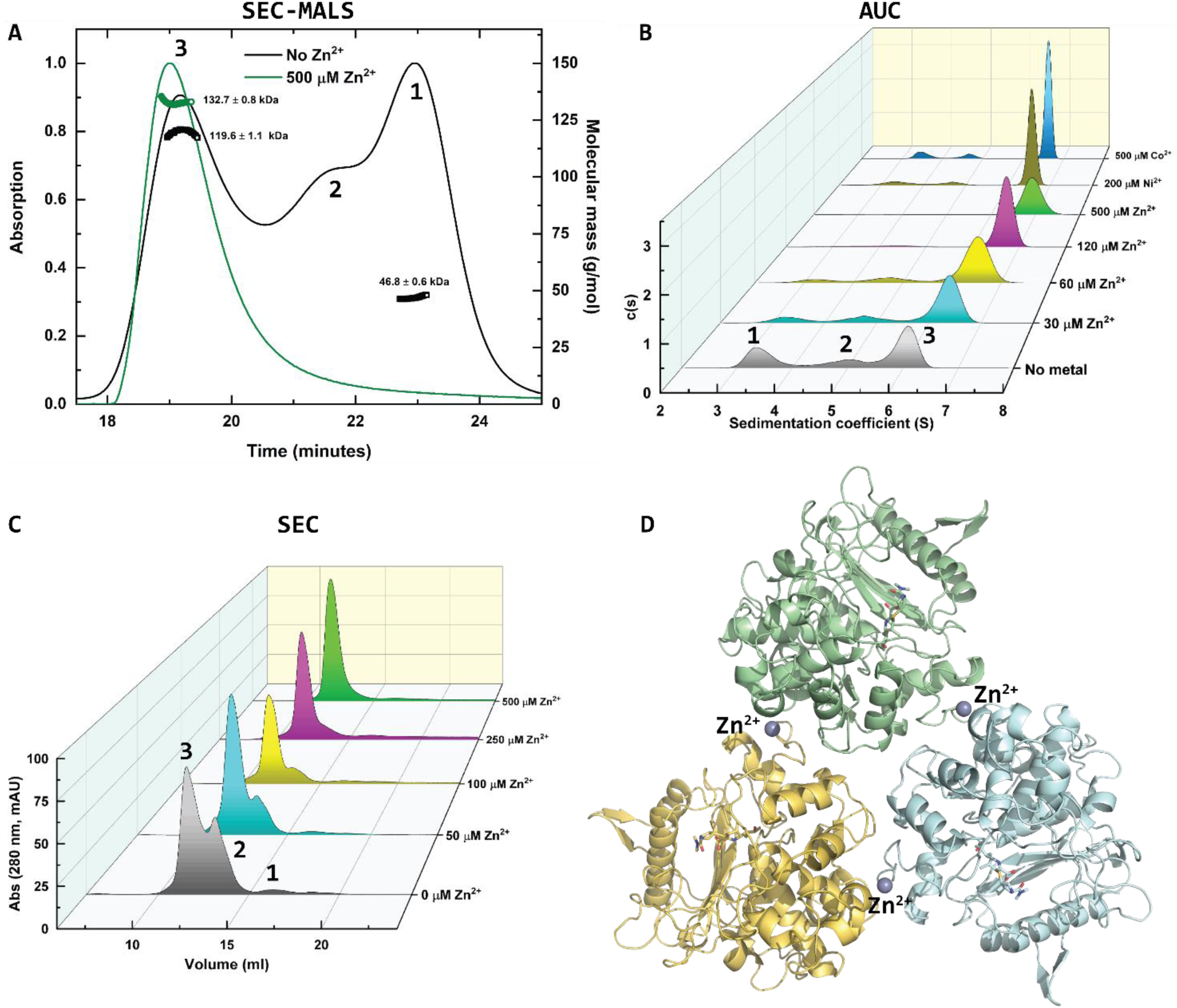
Oligomerization state of PIB-1 in the absence and in the presence of Zn^2+^. **A** Oligomerization state analyzed by SEC-MALS. The black line corresponds to the elution profile of PIB-1 in the absence of Zn^2+^. The green line corresponds to the elution profile of PIB-1 in the presence of 500 μM Zn^2+^. The calculated molecular mass for each elution peak is shown as short lines close to the peak’s maxima. **B** Oligomerization state analyzed by sedimentation velocity (AUC). The metal and the concentration added for each curve is shown on the z axis. **C** Gel filtration chromatography (SEC) of PIB-1 in the absence (gray) and in the presence of increasing amounts of Zn^2+^ (50 μM, cyan; 100 μM, yellow; 250 μM, purple and 500 μM, green). **D** Structure of the PIB-1 trimer showing the three atoms of Zn^2+^ that contribute to the generation of the oligomer.

### Metal binding to PIB-1

We carried out measurements of the dissociation constant (K_d_) for the binding of different metals to PIB-1 using isothermal titration calorimetry (ITC), quenching of the intrinsic protein fluorescence and circular dichroism (CD). The ITC measurements shows that PIB-1 is able to bind Zn^2+^, Ni^2+^ and Co^2+^ with decreasing affinity (Fig 7A-7C). These metals also produce a quenching effect on the intrinsic protein fluorescence (Fig 7D and S3 Fig). Zn^2+^ produce a progressive quenching up to a concentration of 2 mM (Fig 7D, black symbols and S3A Fig), further increasing the concentration of Zn^2+^ produce a milky solution that did not increase the quenching effect (Fig 7D, blue symbols and S3A Fig). This effect does not precipitate the protein and, most likely, is due to the appearance of protein aggregates, as has been reported for other proteins (Mu *et al*, 2011; Tanaka *et al*, 2004). The quenching effect produced by Ni^2+^ saturates at a concentration of 2 mM (Fig 7D, red symbols and S3B Fig), and Co^2+^ produced a quenching effect similar to a collisional quenching, as that caused by potassium iodide (Fig 7D, green symbols, S3C and D Fig). In the case of the last two metals, the solution remains clear up to the maximum concentration used (6 mM). We also measured the effect of the metals on the near-UV circular dichroism spectra of PIB-1. These three metals produce a significant decrease in the intensity of the 250 nm relative maximum (Fig 7E-7G). The intensity of this maximum has been used to calculate the K_d_ for the binding of these metals to PIB-1 (Fig 7E-7G, insets). There is a small effect on the 283 nm maximum that does not seem to be concentration dependent, and over a concentration of 238 μM, the region between 300 and 340 nm that includes a relative minimum at around 303 nm increases its intensity (Fig 7E-7G). The change of the CD signal at the 300 to 340 nm region that occurs with the three metals might indicate an induction of non-specific aggregation, as indicated by the milky aspect of the solution in the presence of high concentrations of Zn^2+^. The K_d_ for the binding of metals to PIB-1 are summarized in Table 2. The non-specific aggregation effect, observed by the milky aspect of the solution and by the change of the 300 to 400 nm region of the near-UV CD spectra at higher concentration of the metals, is reversible. Incubating PIB-1 with a concentration of 796 μM Zn^2+^ produced a milky solution that upon dilution to specific concentrations of the metal reversed the transparency of the solution, and the near-UV CD spectra showed a spectrum similar to that of the final metal concentration without any prior incubation with high concentrations (S4 Fig).

**Fig 7.**
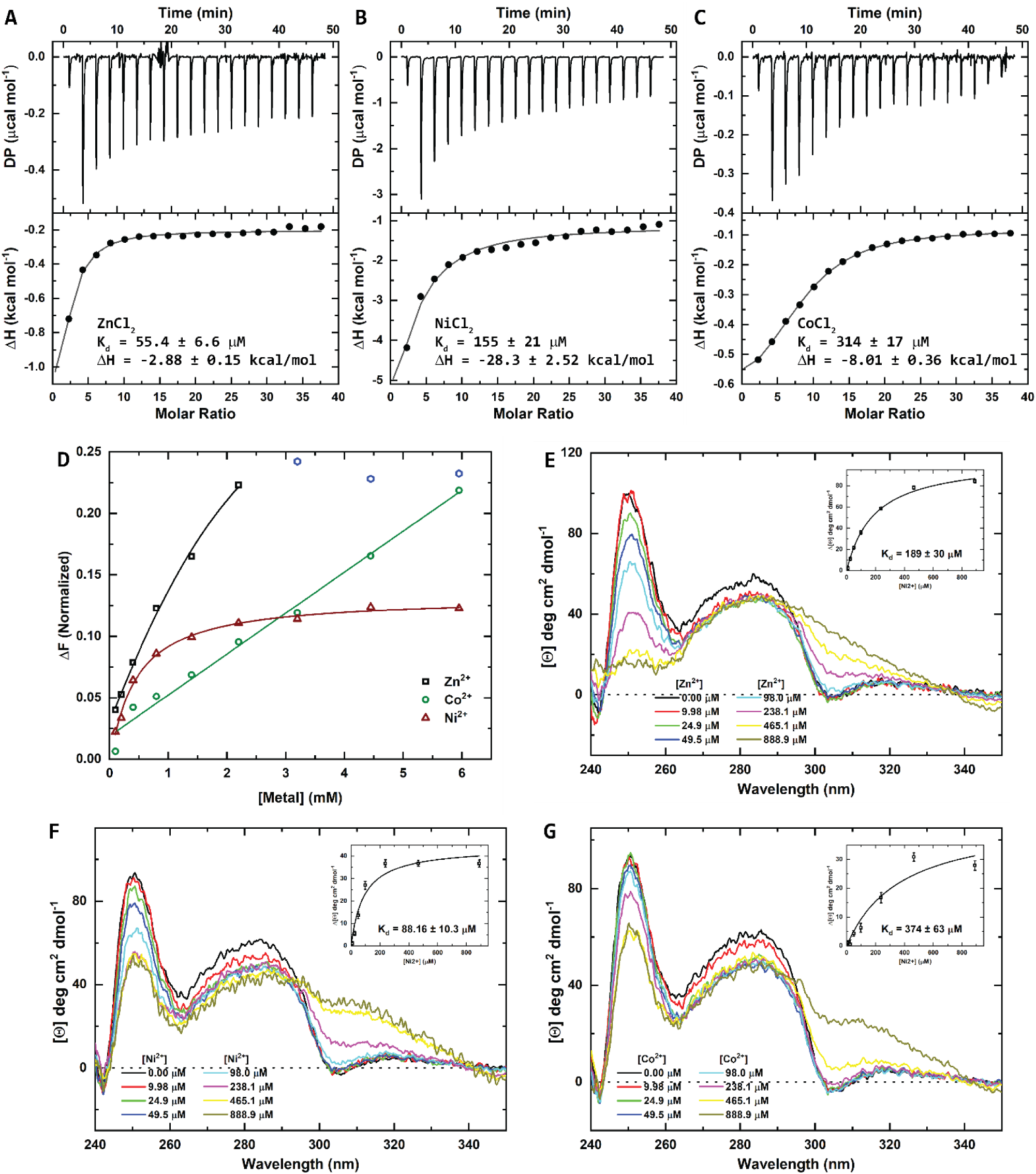
Binding of metals to PIB-1. **A** ITC profile of the binding of Zn^2+^ to PIB-1. **B** ITC profile of the binding of Ni^2+^ to PIB-1. **C** ITC profile of the binding of Co^2+^ to PIB-1. **D** Titrations of the intrinsic protein fluorescence quenching effect of metals to PIB-1. **E** Near-UV circular dichroism spectra of the effect of Zn^2+^ upon binding to PIB-1. The inset shows the titration at 250 nm. **F** Near-UV circular dichroism spectra of the effect of Ni^2+^ upon binding to PIB-1. The inset shows the titration at 250 nm. **G** Near-UV circular dichroism spectra of the effect of Co^2+^ upon binding to PIB-1. The inset shows the titration at 250 nm.

**Table 2.** Dissociation constant of metals for the binding to PIB-1.

| Metal | $K_d$ ($\mu M$) | |
| --- | --- | --- |
|  | Circular dichroism | ITC |
| $Zn^{2+}$ | $189 \pm 30$ | $55.4 \pm 6.6$ |
| $Ni^{2+}$ | $88.2 \pm 10.3$ | $155 \pm 21$ |
| $Co^{2+}$ | $374 \pm 63$ | $314 \pm 17$ |

Altogether, our data show that there is a specific binding of metals by PIB-1, with the metal binding site being formed by residues from two monomers. The binding of the metal drives the oligomerization equilibrium from a state where monomers, dimers and trimers exist towards the formation of trimers.

### Binding of carbapenem antibiotics to PIB-1

The binding of the carbapenems imipenem and meropenem to PIB-1 has been measured by three different techniques: intrinsic protein fluorescence quenching, CD and ITC. The sequential addition of the carbapenem antibiotics produces a strong quenching effect of the intrinsic protein fluorescence (Fig 8A, inset, S5A and S5B Fig). At the highest concentration used (396 μM) the fluorescence signal nearly disappeared. The titrations followed by CD are shown in Fig 8B. The dichroic signal in the near-UV region also was reduced by the addition of the antibiotics. The ITC titrations are shown in Fig 8C and 8D. The binding of imipenem to PIB-1 was slightly endothermic in the absence of metals (Fig 8C) while the presence of Zn^2+^ modified the reaction to be slightly exothermic (Fig 8D). The binding of imipenem in the presence of Ni^2+^ or Co^2+^ was slightly endothermic (S6 Fig). The K_d_ for the two different antibiotics measured using the three independent techniques are very similar (Table 3), indicating that both antibiotics might bind similarly to PIB-1. Also, the presence of any of the three metals did not produce significant differences in the values of K_d_. This, most likely, indicates that the metal is not involved in the binding of the antibiotics and do not bind to the active site, as seen by X-ray crystallography.

**Fig 8.**
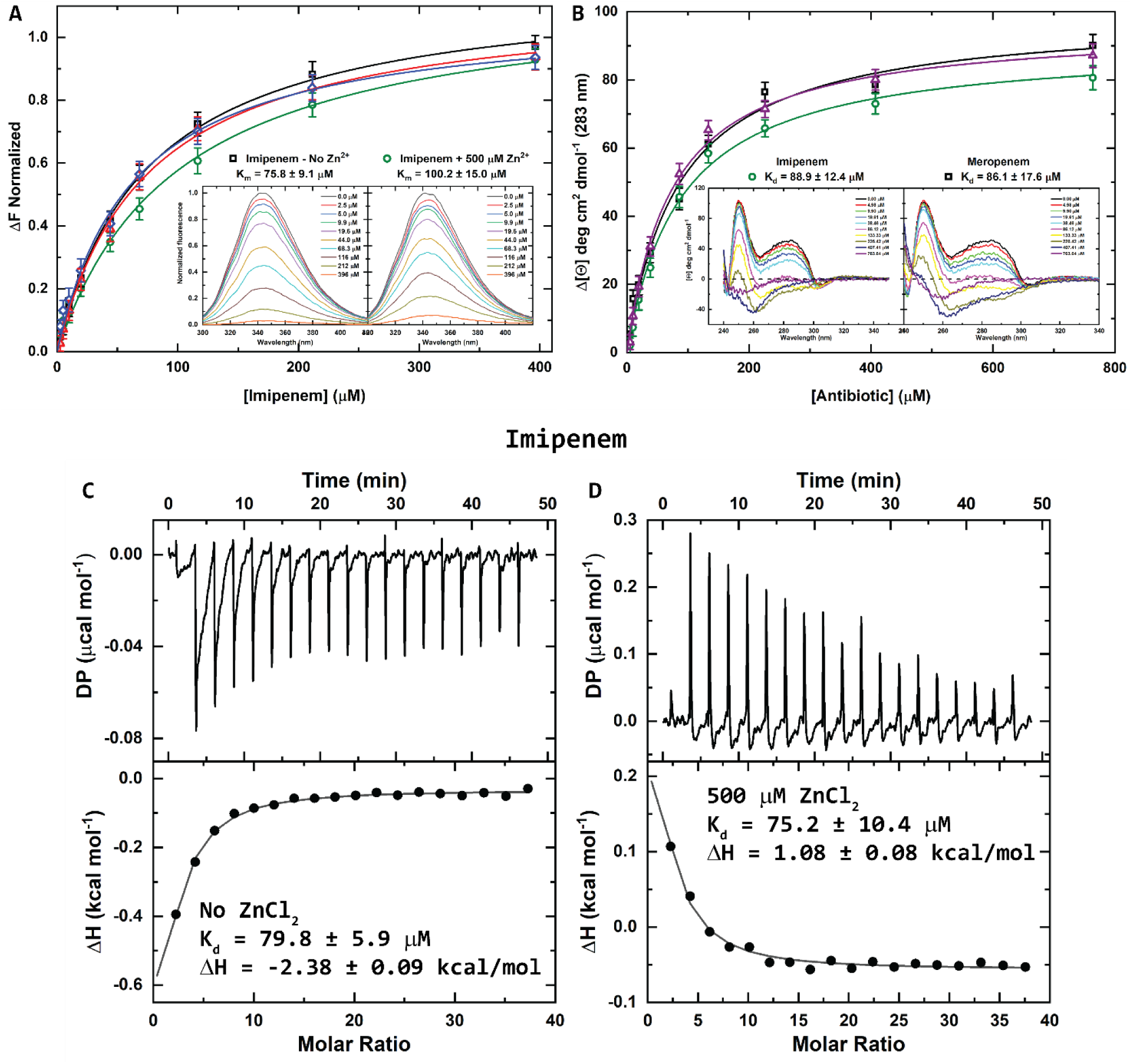
Binding of imipenem and meropenem to PIB-1. **A** Titrations of the intrinsic protein fluorescence quenching of PIB-1 upon binding of Imipenem in the absence (black squares) and in the presence (green circles) of Zn^2+^. The inset shows the fluorescence spectra for the titration. **B** Titrations of the near-UV circular dichroism of PIB-1, at 283 nm, upon binding of Imipenem in the absence (black squares) and in the presence (purple triangles) of Zn^2+^, and upon binding of meropenem in the absence of Zn^2+^ (green circles). The inset shows the Near-UV circular dichroism spectra for the titration. **C** ITC profile of the binding of imipenem to PIB-1 in the absence of metals. **D** ITC profile of the binding of imipenem to PIB-1 in the presence of Zn^2+^. The experiments were performed in duplicate and the error bars represent the standard deviation.

**Table 3.** Dissociation constant of meropenem and imipenem for the binding to PIB-1.

| | $K_d$ ($\mu\text{M}$) | | |
| --- | --- | --- | --- |
|  | Florescence quenching | Circular dichroism | ITC |
| Imipenem | $75.8 \pm 9.1$ | $88.8 \pm 12.4$ | $79.8 \pm 5.9$ |
| Imipenem + 500 $\mu\text{M}$ $\text{Zn}^{2+}$ | $100.2 \pm 15.0$ | $74.2 \pm 6.2$ | $75.2 \pm 10.4$ |
| Imipenem + 500 $\mu\text{M}$ $\text{Ni}^{2+}$ | n.d. | n.d. | $90.1 \pm 7.3$ |
| Imipenem + 500 $\mu\text{M}$ $\text{Co}^{2+}$ | n.d. | n.d. | $86.4 \pm 8.2$ |
| Meropenem | $74.8 \pm 10.3$ | $86.1 \pm 17.6$ | n.d. |
| Meropenem + 500 $\mu\text{M}$ $\text{Zn}^{2+}$ | $63.2 \pm 8.8$ | n.d. | n.d. |

n.d. = not determined.

### Kinetic measurements

PIB-1 is able to catalyze the opening of the β-lactam ring of nitrocefin, a chromogenic β-lactamase substrate, with very low efficiency (Fig 9A, black symbols). The presence of metals (Zn^2+^, Ni^2+^, Co^2+^) increases the catalytic constant (k_cat_) with a slight decrease of the Michaelis-Menten constant (K_m_) for the catalysis of nitrocefin that results in an increase in the catalytic efficiency of more than 14-fold (Table 4). It is also able to catalyze the opening of the β-lactam ring of imipenem and meropenem in the presence of Zn^2+^ (Fig 9B and S7A Fig), as seen by X-ray crystallography for meropenem (Fig 1E). In the absence of metals, it has a very poor capacity for the catalysis of the carbapenems, but it is enhanced by the presence of these metals (Fig 9B and S7A Fig), as reported previously for Zn^2+^ (Fajardo *et al*., 2014). The catalysis of the imipenem and meropenem by PIB-1 does not follow a Michaelis-Menten kinetics (Fig 9B and S7A Fig). Imipenem and meropenem are able to inhibit the catalysis of nitrocefin by PIB-1 (Fig 9C and S7B Fig).

**Fig 9.**
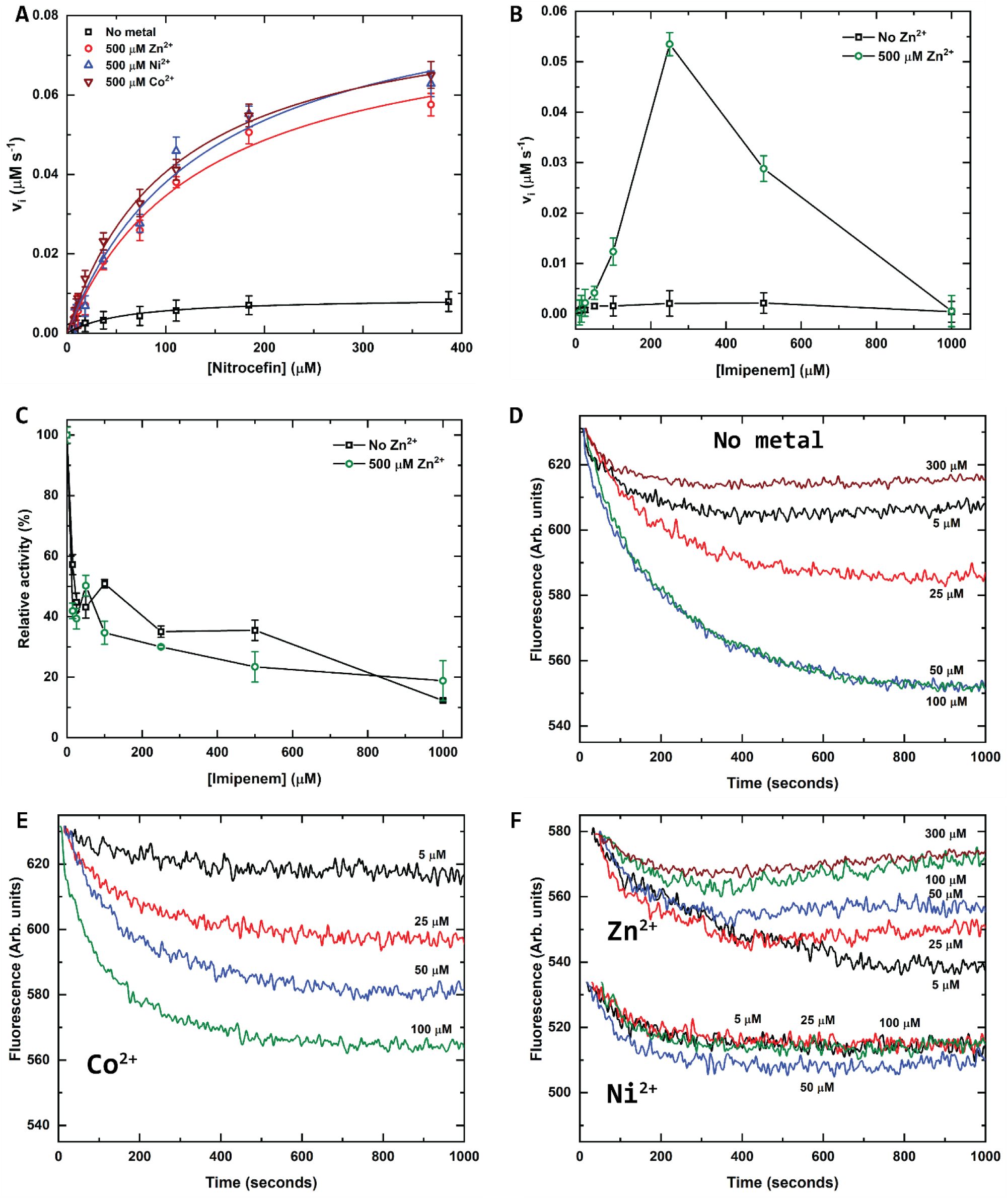
Kinetic measurements of the catalysis of antibiotics by PIB-1. **A** Steady-state kinetics of the catalysis of nitrocefin by PIB-1 in the absence (black symbols) and in the presence of 500 μM of Zn^2+^ (red symbols), Ni^2+^ (blue symbols) or Co^2+^ (brown symbols) followed by the absorption change at 486 nm. **B** Steady-state kinetics of the catalysis of imipenem by PIB-1 in the absence (black symbols) and in the presence of 500 μM of Zn^2+^ (green symbols) followed by the absorption change at 298 nm. **C** Imipenem inhibition of the catalysis of nitrocefin by PIB-1 in the absence (black symbols) and in the presence of 500 μM of Zn^2+^ (green symbols) followed by the absorption change at 486 nm. **D** Kinetics of the binding of imipenem to PIB-1 in the absence of metals followed by the quenching effect of the antibiotic on the intrinsic protein fluorescence. The origin points of the curves have been shifted to coincide with that of the minimum concentration of antibiotic. **E** Kinetics of the binding of imipenem to PIB-1 in the presence of 500 μM Co^2+^ followed by the quenching effect of the antibiotic on the intrinsic protein fluorescence. **F** Kinetics of the binding of imipenem to PIB-1 in the presence of 500 μM Zn^2+^ or Ni^2+^ followed by the quenching effect of the antibiotic on the intrinsic protein fluorescence. The curves representing the effect of Ni^2+^ have been shifted down for clarity. The experiments were performed in duplicate and the error bars represent the standard deviation.

**Table 4.** Steady-state kinetic parameters for the hydrolysis of Nitrocefin by PIB-1.

| | $K_m$ ($\mu\text{M}$) | $k_{\text{cat}}$ ($\text{s}^{-1}$) | $k_{\text{cat}}/K_m$ ($\text{s}^{-1} \mu\text{M}^{-1}$) |
| --- | --- | --- | --- |
| Nitrocefin | $95.3 \pm 8.9$ | $0.0041 \pm 0.0005$ | $43 \times 10^{-5}$ |
| Nitrocefin + 500 $\mu\text{M}$ $\text{Zn}^{2+}$ | $136.2 \pm 12.6$ | $0.0370 \pm 0.0012$ | $600 \times 10^{-5}$ |
| Nitrocefin + 500 $\mu\text{M}$ $\text{Ni}^{2+}$ | $139.6 \pm 10.1$ | $0.0412 \pm 0.0015$ | $666 \times 10^{-5}$ |
| Nitrocefin + 500 $\mu\text{M}$ $\text{Co}^{2+}$ | $107.7 \pm 15.3$ | $0.0381 \pm 0.0017$ | $783 \times 10^{-5}$ |

The kinetics for the binding of the antibiotics to PIB-1 in the presence and in the absence of metals are shown in Fig 9D-9F, followed by quenching of the intrinsic protein fluorescence. The majority of the quenching effect occurs within the first few seconds (<10 sec). Then, there is a gradual change during the following 5 to 10 minutes, indicating some slow process. This slow change represents a 6.6 ± 3.5% of the total fluorescence signal decrease due to the binding of the antibiotic. The presence of metals does not eliminate this slow fluorescence signal change, instead Zn^2+^ and Ni^2+^ reduced this signal change to 3.3 ± 2.4% and 2.9 ± 0.4%, respectively (Fig 9F). This, most likely, indicates that the oligomer induced by the presence of these metals is able to bind the antibiotics slightly more efficiently than the oligomerization mixture we have in their absence. Co^2+^ does not decrease the quenching signal change as the other two metals, 5.6 ± 4.1% (Fig 9E). This might be due to the lower affinity of this metal (Fig 6C and 6G, Table 2). Once reached the final value, the fluorescence signal does not change for the next hour (S8 Fig), which likely indicates that the hydrolyzed antibiotics remain bound to the protein, acting at this stage more as inhibitors than as substrates. The enzymatic activity of PIB-1 can be recovered after incubation with imipenem. When PIB-1 is incubated with any of the three metals and the concentration of imipenem is diluted to a low concentration (from 500 to 5 μM) the activity of the protein, using nitrocefin as the substrate, PIB-1 is able to degrade the substrate (S9 Fig). When we maintain the inhibitory concentration of imipenem (500 μM), the nitrocefin hydrolyzing activity is not recover (S9 Fig). As seen previously with the susceptibility experiments (Table 1) PIB-1 is not able to catalyze the degradation of cephalosporins (cefepime, cefotaxime and ceftazidime, S10 Fig). S11 Fig shows the molecular formula of the antibiotics used in these kinetic studies.

The presence of metals increases the enzymatic activity of PIB-1, most likely, by shifting the oligomerization equilibrium towards the trimers. PIB-1 is able to catalyze the degradation of imipenem and meropenem and it does not seem to follow a Michaelis-Menten kinetic model. Our data indicate that the hydrolyzed antibiotic remains bound to the protein ant its release is slow, suggesting that the product release is the slower step of the reaction. However, the hydrolyzed antibiotic can be forced out by another substrate (as seen with nitrocefin). We hypothesize that this β-lactamase might be the product of an adaptation to the presence of high concentration of Zn^2+^ that can be found in hospital environments. In fact, the presence of concentrations of around 560 μM Zn^2+^ in the eluates from siliconized latex urinary catheters induce resistance to imipenem in *P. aeruginosa* has been reported (Conejo *et al*, 2003; Martínez-Martínez *et al*, 1999). This concentration of Zn^2+^ is between 5-10 times that of the K_d_ of PIB-1 and might be possible that it contributes to the resistance of *P. aeruginosa* to carbapenems in this environment.

## Discussion

AmpC β-lactamases constitute a very diverse family of antibiotic resistance determinants whose primary structure can be conserved by just 20 % but they present a well conserved tertiary structure. In particular, three motifs and 70 amino-acids are conserved on more than 90 % of the so far described AmpC β-lactamases (Philippon *et al*., 2022). While their substrate range is variable, all of them share a favorite type of substrate: cephalosporins, and none of them efficiently hydrolyze carbapenems. Herein, we describe the crystal structure of the β-lactamase PIB-1 from *P. aeruginosa*. Despite the primary structure of this enzyme being dissimilar to other AmpC β-lactamases, containing insertions and deletions absent in other members of the AmpC family, its crystal structure fits within the AmpC category. Noteworthy, PIB-1 only presents one of the three conserved catalytic motifs and 24 of the 70 amino acids that are highly conserved on more than 90 % of the described AmpC β-lactamases (Philippon *et al*., 2022). In agreement with these findings, phylogenetic analysis shows that PIB-1 forms an outgroup within the AmpC family of β-lactamases. In addition, and differing with what has been described for other AmpC β-lactamases, PIB-1 does not contribute to resistance neither to cephalosporins nor to any other β-lactams besides carbapenems. Moreover, biochemical analysis supports that it does not degrade cephalosporins, while able of degrading carbapenems. Also, the activity of the protein depends on the presence of metals. Based in its substrate profile and metal dependence, PIB-1 should be classified as a B-type β-lactamase. However, the structural analysis here presented shows unequivocally that it is an AmpC β-lactamase. It is worth mentioning that, while metals bind the catalytic domain of B-type β-lactamases, their role for increasing PIB-1 activity is completely different. In this case, they do not bind to the catalytic domain, but promote the formation of trimers, likely more active that dimers and monomers for hydrolyzing carbapenems. The differential structural features of PIB-1, including the lack of several conserved amino-acids, might be behind this novel substrate profile in this new type of AmpC β-lactamase. Databases of antibiotic resistance determinants are strongly biased in favor of bacterial pathogens and, eventually, commensals. Consequently, although the sequences of nearly 5000 AmpC β-lactamases have been reported until now, most of them belong to *Enterobacteriaceae* and *Pseudomonadaceae*. The fact that PIB-1 is an AmpC β-lactamase that does not contain most of the highly conserved residues of this category and that its substrate profile and metal dependence resemble the ones of B-type β-lactamases indicates that the group of AmpC β-lactamases might be even more diverse than expected, containing enzymes with substrate profiles and primary structures that differ to those of classical -described-AmpC β-lactamases.

## Materials and Methods

### Protein Expression and Purification

The expression and the purification of the native protein was carried out as described previously (Fajardo *et al*., 2014). The expression of the selenomethionine (SeMet)-derivatized protein was carried out according to a modification of the inhibition of methionine-biosynthesis pathway technique (Van Duyne *et al*, 1993) as described (Spinola-Amilibia *et al*, 2016). The SeMet-derivatized protein was purified as the native protein.

### Crystallization and Data Collection

Crystallization trials were performed at 295 K using the sitting-drop vapor-diffusion method with commercial screening solutions including JBScreen Classic and Wizard I–IV (Jena Bioscience, Jena, Germany) in 96-well sitting-drop plates (Swissci MRC; Molecular Dimensions, Suffolk, England). Drops were set up by mixing equal volumes (0.2 μL) of a protein solution (10 mg mL^−1^) and reservoir solution using a Cartesian Honeybee System (Genomic Solutions, Irvine, CA, USA) nano-dispenser robot and equilibrated against 50 μL of reservoir solution. Three different protein solutions were used in the crystallization assays: protein alone, protein with 2 mM ZnCl_2_, and protein with 2 mM ZnCl_2_ and 2 mM meropenem. Crystals of the protein alone appeared after three days in 0.1 M TRIS-HCl buffer at pH 8.6 containing 17.5% PEG 4000 and 0.2 M CaCl_2_. Crystals of the protein in complex with meropenem appeared after one week in 0.1 M sodium cacodylate buffer at pH 6.5 containing 20% PEG 1000 and 0.2 M MgCl_2_. Crystals of the SeMet-derivatized protein appeared after two days in 0.1 M sodium acetate buffer at pH 4.6 containing 8% PEG 4000. Crystals of the native protein and the meropenem complex were harvested, for data collection, in paratone and flash-cooled in liquid nitrogen. Crystals of the SeMet-derivatized protein were harvested in the crystallization solution containing 30% PEG 400 and flash-cooled in liquid nitrogen. X-Ray data collection experiments were performed at the ALBA Synchrotron (Cerdanyola del Vallès, Spain) BL13 XALOC beamline and at the European Synchrotron Radiation Facility (ESRF, Grenoble, France) ID29 beamline. Data were indexed and integrated, scaled and merged using XDS (Kabsch, 2010).

### Structure Determination

The structure could not be solved by molecular replacement, despite the presence of similar proteins in the PDB. SAD data from crystals of the SeMet-derivatized protein, measured at the peak of the selenium, were used to solve the structure. This protein was solved using the program CRANK-2 (Ness *et al*, 2004) implemented in the CCP4 suite (Winn *et al*., 2011). The final model comprised 1542 residues in 42 fragments that represent 81.2% of the sequence, the Figure of Merit (FOM) is 0.8279 after 10 cycles of model building with a R_factor_ of 0.2926 and a R_free_ of 0.3297. A final macromolecular reciprocal space refinement lowered the value of R_factor_ to 0.2784 and of R_free_ to 0.3147. This model was manually built with Coot (Emsley *et al*, 2010) and refined with Phenix (Liebschner *et al*, 2019). The structure of the SeMet-protein was used to solve the structure of the PIB-1-meropenem complex by molecular replacement with Phaser (McCoy *et al*, 2007). The structure of the apo protein was solved using the structure of the PIB-1-meropenem complex. The initial models were first refined using Phenix (Liebschner *et al*., 2019) and alternating manual building with Coot (Emsley *et al*., 2010). The final models were obtained by repetitive cycles of refinement; solvent molecules were added automatically and inspected visually for chemically plausible positions. The meropenem molecules were added manually. The stereochemical quality of the models were assessed with MolProbity (Chen *et al*, 2010). The structural figures were generated using the <u>PyMol</u> program (www.pymol.org). Data processing and refinement statistics are listed in S1 Table.

### Size exclusion chromatography-multi angle light scattering (SEC-MALS)

The experiments were performed using a Superdex 75 10/300 GL column (Cytiva) attached in-line to a DAWN-HELEOS light scattering detector and, in parallel, to an Optilab rEX refractive index detector (Wyatt Technology). Protein samples (200 μL) with a concentration of 1 mg/mL were injected into the size exclusion column equilibrated with 20 mM Tris-HCL pH 8.0 and 100 mM NaCl, in the absence and in the presence of 500 μM ZnCl_2_, and the chromatography process was carried out at room temperature. Data acquisition and analysis were performed using ASTRA software (Wyatt) to calculate the molecular weight (MW) of the protein species that eluted from the column.

### Size exclusion chromatography (SEC)

The protein was chromatographed on a Enrich SEC 650 10 x 300 column (Bio-Rad) using an ÄKTApurifier system (Cytiva). Aliquots of 100 μL of PIB-1 at a concentration of 1 mg/mL in 10 mM TRIS-HCl buffer at pH 8.0 and 100 mM NaCl in the absence and in the presence of metals were injected into the system and run with a flow rate of 1.0 mL/min at 4 °C. Protein elution was recorded at 280 nm. The buffer was supplemented with the desired concentration of metal (ZnCl_2_, NiCl_2_ and CoCl_2_) for the chromatography experiments in the presence of metals. The column was calibrated with the following molecular weight markers: thyroglobulin (Mr = 669 kDa), ferritin (Mr = 440 kDa), catalase (Mr = 232 kDa), aldolase (Mr = 158kDa), bovine serum albumin (Mr = 67 kDa), lysozyme (Mr = 14 kDa) and fluorescein (Mr = 0,33 kDa).

### Analytical ultracentrifugation (sedimentation velocity)

Sedimentation velocity experiments were carried out using an Optima XL-I analytical ultracentrifuge (Beckman-Coulter) equipped with UV-visible absorbance optics at 20 °C in an AnTi50 rotor and 12-mm double sector Epon-charcoal centerpieces. Measurements were performed at 48,000 rpm, registering successive entries every minute at 280 nm. The protein was at a concentration of 0.4 mg/mL in 10 mM TRIS-HCl buffer at pH 8.0 and 100 mM NaCl in the absence and in the presence of ZnCl_2_, NiCl_2_ and CoCl_2_ at the desired concentrations. The sedimentation coefficient distributions were calculated by least squares boundary modeling of sedimentation velocity data using the c(s) method(Dam & Schuck, 2005; Reynolds & McCaslin, 1985; Schuck, 2000), as implemented in the SEDFIT software (Version 16.1c). Partial specific volume, v, for PIB-1 was calculated at 0.731 mg/mL based on the amino acid sequence.

### Dynamic light scattering (DLS)

Dynamic light scattering experiments were performed with a DynaPro instrument (Protein Solutions Inc.). Measurements were taken at 20 °C. Aliquots of the samples used for the sedimentation velocity experiment was used to measure the DLS. The software provided by the manufacturer was used to calculate the diffusion coefficient of the protein.

### Isothermal Titration Calorimetry (ITC)

ITC experiments were performed using a PEAQ-ITC calorimeter (Malvern, Westborough, MA, USA). Injections of 2 μL of metal or imipenem solutions were added at 150 s intervals (up to adding 36.4 μL) at 25 °C with stirring at 750 rpm. The concentration of PIB-1 in the cell was 25.4 μM, the imipenem concentration in the syringe was 5 mM and the concentration of the metals in the syringe was 5 mM. The concentration of the metal in the titrations to measure the binding of imipenem in the presence them was 500 μM in the reservoir and in the syringe. Data processing was performed using the MiroCal PEAQ-ITC Analysis software. In each titration, a fitted offset parameter was applied to account for potential background. The experimental data were fitted to a theoretical titration curve with ΔH (binding enthalpy in kcal mol^-1^) and K_d_ (dissociation constant) as adjustable parameters. Due to the low c value (0.08 – 0.45) in all experiments performed the value of n (number of binding sites per monomer) was fixed at 1, because the low reliability in the determination of this parameter in these conditions(Turnbull & Daranas, 2003). This value was taken from the information provided by the X-ray crystal structure that indicates the binding of one molecule of antibiotic to the active site of PIB-1 and one atom of Zn^2+^ per monomer.

### Enzyme kinetics

A continuous spectrophotometric assay was used for the activity studies. Data were collected using a Cary 4000 spectrophotometer (Varian) equipped with a Peltier temperature control and a kinetics software package. The absorbance was monitored at the following wavelengths: nitrocefin (486 nm and Δε = 20,500 M^-1^ cm^-1^), imipenem (297 nm and Δε = 10,930 M^-1^ cm^-1^), meropenem (298 nm and Δε = 7,200 M^-1^ cm^-1^), cefepime (260 nm and Δε = 750 M^-1^ cm^-1^), cefotaxime (260 nm and Δε = 1830 M^-1^ cm^-1^) and ceftazidime (260 nm and Δε = 8860 M^-1^ cm^-1^). The initial velocities (v_i_) were determined from the linear phase of the reaction time courses. The observed velocities were plotted as a function of the antibiotic concentration and fit non-linearly to the Michaelis-Menten equation using the program GraphPad Prism 9.5.5 to obtain the steady-state kinetic parameters V_max_ and K_m_. Reactions were carried out at 37 °C in 10 mM Na-HEPES at pH 7.5, 100 mM NaCl, and varying concentrations of the β-lactam substrate were initiated by the addition of enzyme. The concentration of protein used was 0.1 mg/mL (2.21 μM). Figures were generated with the program ORIGIN 2018 (www.originlab.com).

### Recovery of the enzymatic activity

The recovery of the enzymatic activity after incubation with imipenem was measured by a continuous spectrophotometric assay using nitrocefin as the substrate. The protein at a concentration of 11 mg/mL (253 μM) was incubated for 15 minutes at room temperature in the presence of 500 μM imipenem and 500 μM of metal, then it was diluted with a solution containing 362 μM nitrocefin in the absence or in the presence of 500 mM of ZnCl_2_, NiCl_2_, or CoCl_2_, and the enzymatic activity was recorded.

### Inhibition of the activity of PIB-1 on nitrocefin by imipenem and meropenem

The inhibition of the enzymatic activity of PIB-1 using nitrocefin as the substrate by imipenem and meropenem was carried out by mixing the protein with the substrate (nitrocefin) and the antibiotics, and the activity was followed spectrophotometrically at 486 nm.

### Fluorescence measurements

Fluorescence was measured using the Luminescence Spectrometer L 50 B from Perkin Elmer. Spectra were recorded with an excitation wavelength of 280 nm using a 5 x 10 mm cell at 25 °C with excitation and emission slits of 3 and 5 nm, respectively. Measurements were carried out at a protein concentration of 0.062 mg/mL in 10 mM Tris-HCl buffer at pH 8.0. The spectra were corrected for the buffer and ligand contribution, and for the dilution factor due to the addition of the ligand. Equilibrium K_d_ values were determined by fitting the data to a nonlinear regression model for one site-specific binding using GraphPad Prism 9.5.5, assuming one binding site. Figures were generated with the program ORIGIN 2018 (www.originlab.com).

### Time course of the quenching effect

The time course of the quenching effect was measured using the intrinsic protein fluorescence. For the short-time courses (16 minutes) the protein was mixed with the metal and the antibiotic and the data were collected immediately. For the long-time courses (60 minutes) the protein was mixed with the metal and the antibiotic, incubated for 15 minutes and then the data were collected immediately.

### Circular dichroism measurements

Circular dichroism measurements were carried out on a JASCO J-720 (Jasco, Tokyo, Japan) spectropolarimeter equipped with a Peltier type temperature controller and a thermostatized cuvette cell linked to a thermostatic bath. Near-UV spectra were recorded in 1 cm path length quartz cells at 25 °C with a response time of 1 sec and a band width of 1 nm, 10 consecutive scans were collected and averaged. The spectra were corrected for the contribution of the buffer and the ligands, and by the dilution factor due to the addition of the ligand. The protein concentration used was 1.0 mg/mL in 10 mM Tris-HCl buffer at pH 8.0. The observed ellipticities were converted into the molar ellipticities [θ] based on a mean molecular mass per residue of 111.51 Da. Figures were generated with the program ORIGIN 2018 (www.originlab.com).

### Reversibility of the binding of metals

The reversibility of the aggregation effect produced by Zn^2+^ was carried out by near-UV circular dichroism. A 20 μL solution of PIB-1 at a concentration of 448 μM was incubated in the presence of a concentration of Zn^2+^ of 796 μM for 15 minutes, then the solution was diluted 25-fold to 500 μL. This solution was diluted in the absence of Zn^2+^ or in the presence of several concentrations of the metal, and the near-UV CD spectra were collected.

### Bacterial strains and determination of susceptibility to antibiotics

Bacterial strains and plasmids used in this work are listed in Supplementary Table 5. Bacteria were grown in glass tubes in Lysogeny Broth (LB) (Lenox, Pronadisa) with shaking at 250 rpm at 37 °C. MICs of β-lactams antibiotics were determined at 37 °C, in Mueller Hinton (MH), using E-test strips (MIC Test Strip, Liofilchem®) in the presence of 1 mM IPTG and 1 mM ZnSO_4_.

## Acknowledgments

This research was funded by the MCIN/AEI /10.13039/501100011033, grant PID2020-113521RB-I00 (to JLM) and by the Consejo Superior de Investigaciones Científicas, PIE-202020E224 (to AR). The authors wish to thank the staff of beamlines ID29 (ESRF Synchrotron) and BL13-XALOC (ALBA Synchrotron) for their generous and much appreciated support and the CIB Molecular Interaction Service for the SEC-MALS analysis and the sedimentation velocity experiments.

